# ω-imidazolyl-alkyl derivatives as new preclinical drug candidates for NASH therapy

**DOI:** 10.1101/685115

**Authors:** Torsten Diesinger, Alfred Lautwein, Vyacheslav Buko, Elena Belonovskaya, Oksana Lukivskaya, Elena Naruta, Siarhei Kirko, Viktor Andreev, Radovan Dvorsky, Dominik Buckert, Sebastian Bergler, Christian Renz, Dieter Müller-Enoch, Thomas Wirth, Thomas Haehner

## Abstract

Cytochrome P450 2E1 (CYP2E1) and its production of ROS play an essential role in the development and progression of inflammatory liver diseases such as alcoholic steatohepatitis. For this isoenzyme we have developed two new inhibitors - 12-imidazolyl-1-dodecanol (I-ol) and 1-imidazolyldodecane (I-an) - and wanted to test their effect on the related disease of non-alcoholic steatohepatitis (NASH). The fat-rich Lieber-DeCarli diet, which was administered over the entire experimental period of 16 weeks, was used for disease induction in the rat model, while the experimental substances were administered in parallel over the last four weeks. This high-calorie diet pathologically altered the ROS balance, the amount of adipocytokines, TNF-α and lipids as well as the activities of liver enzymes. Together with the histological examinations, the conclusion could be drawn that the diet led to the formation of NASH. I-ol and to a lesser extent I-an were able to shift the pathological values towards the normal range - despite continued administration of the noxious agent. I-ol, in particular, showed an extremely good tolerability in the acute toxicity study in rats. Thus, CYP2E1 appears to be a suitable drug target as well as I-ol and I-an promising drug candidates for the treatment of NASH.

## Introduction

From a historical point of view, NASH was defined by a similar result in the histological examination of liver biopsies of patients as in alcoholic steatohepatitis (ASH), whereas an influence of alcohol had been excluded(1). Even today, liver biopsy is the only reliable proof for the diagnosis of non-alcoholic steatohepatitis (NASH), provided that alcohol, a viral origin, toxins or an autoimmune background can be excluded.

NASH can be a terminal liver disease. It belongs to the spectrum of non-alcoholic fatty liver disease (NAFLD), which also includes liver steatosis, liver fibrosis, liver cirrhosis and hepatocellular carcinoma.

However, there are sometimes inconsistencies in the use of the term NAFLD, because some authors exclude liver steatosis/fatty liver from NAFLD or take both terms as equivalent. The histology of NASH is defined by the simultaneous occurrence of steatosis and inflammation with injury to hepatocytes in the sense of ballooning. Signs of progressive fibrosis are not mandatory. Many intermediate forms between liver steatosis and steatohepatitis seem to complicate this classification. NASH carries the risk of developing hepatocellular carcinoma (HCC) and cholangiocarcinoma. However, HCC does not necessarily arise from NASH via the developmental step of hepatic cirrhosis - unlike ASH, where this is the almost exclusive course of development. Patients diagnosed with NAFLD also have an increased risk of atherosclerosis and thus cardiovascular disease(2).

Previous statistical data on NASH or NAFLD are not very robust and are mostly based on a small sample size. The prevalence of NAFLD has been estimated at 20 % - 30 % in Western countries. A meta-analysis of epidemiological data has recently been published(3), estimating the global presence of NASH at 59 % of biopsy-confirmed patients with NAFLD. On the other hand, it was estimated that 47 % of patients with NASH worldwide suffer from diabetes, 72 % from hyperlipidemia and 68 % from hypertension. In 41 % of patients, NASH leads to liver fibrosis. Approximately 82 % of patients with NASH were found obese in this study. This does not mean that lean people cannot fall ill with NAFLD or NASH. 66 % - 83 % of patients with NAFLD are said to have markers for insulin resistance. This implies an association between NASH, hypertriglyceridemia or dyslipidemia and diabetes mellitus type 2 or insulin resistance (IR), all of which are linked by the so-called metabolic syndrome.

Until nearly a decade ago, NAFLD/NASH was first diagnosed almost exclusively by abdominal sonography and determination of liver enzyme activity in serum, the most prominent of which were alanine aminotransferase (ALT) and aspartate aminotransferase (AST)(4). Today, CT and MRI exami-nations, in particular MR elastography (MRE), are used to characterize even different stages of fibrosis. A large panel of markers for the early detection of NAFLD is discussed(5). These markers include TNF-α, adiponectin, leptin, γ-glutamyltransferase (GGT), ALT and AST. NASH has some of these markers in common with cardiovascular disease, insulin resistance or metabolic syndrome in general.

Results from studies with different rodent models led to the hypothesis that free fatty acids and not a high content of triglycerides in hepatocytes are the true origin of NASH(6). However, high triglyceride levels in liver and serum are still discussed as the main causes of NAFLD (7).

Cyp2e1 plays an important role in the pathogenesis of NASH, where increased hepatic protein expression(8), combined with increased mRNA content(9) and enzymatic activity(10), is observed.

A causal link between NASH and the formation of oxidative stress/ROS stress can be established by CYP2E1 - analogous to the pathogenesis of ASH (11). It has recently been published that insulin resistance (IR) can be linked to CYP2E1 via the anti-apoptotic protein Bax inhibitor-1 (BI-1)(12). This protein occupies an important position in the regulation of CYP2E1 and thus of ER/ROS stress.(13). Furthermore, BI-1 is said to influence the regulation of insulin receptor signaling and thus induce the development of IR when mice receive high-fat diet (HFD). IR is also induced in the liver cell line HepG2 by the saturated fatty acid palmitate via the generation of ROS stress caused by CYP2E1.

There is no single treatment or one approved drug that would be indicated for all four stages of NAFLD or different patient groups. Lifestyle interventions are recommended at an early stage. End-stage liver disease is an indication for liver transplantation, but a definitive cure is only possible in some patients. Almost one third of the patients who received a liver transplant for NASH will have a recurrence of the disease in the trans-planted liver(14). The present status of clinical development with a currently large number of studies was recently published(15).

We have already successfully studied new inhibitors of CYP2E1 in an animal model of alcoholic steatohepatitis. Taking into account the above relationships between CYP2E1, ROS and NASH, we now wanted to test these inhibitors in a model of high-fat diet (HFD) for rats. In addition, we tested the inhibitors in acute toxicity studies with rats according to ICH guidelines.

## Material and Methods

### Animals, diets and treatment

I-ol, I-an and I-phosphocholine were synthesized at the University of Ulm, Germany (see US patent No. US 8153676 B2). Ursodeoxycholic acid (UDCA) was supplied by Prodotti Chimici e Alimentari S.P.A., Basaluzzo, Italy. Hydroxypropylmethylcellulose (Hypromellose, HPMC) was purchased from Shin-Etsu Chemical Co. Ltd, Cellulose Division, Tokyo, Japan. All other chemicals used in this study were of finest analytical grade.

#### (A) NASH study

Female Wistar rats aged 80 - 90 days and weighing 180-200 g were used at the beginning of the experiments. Rats were divided into ten groups based on the diets and experimental substances obtained [Supplement Figure 1]. Group 1 identified the first control group in which a normal standard solid diet was fed. Group 2 as the second control group was characterized by a standard amount of fat content that was part of the liquid ’physiological’ Lieber-DeCarli (LDC) diet. All other groups received the liquid ’disease-causing’ LDC diet [Supplement Table 1], contained a high content of unsaturated fatty acids. Both LDC diets combined eating and drink-ing in liquid form and were produced by ssniff Spezialdiäten GmbH, Soest (Germany). All diets were applied over the entire trial period of 16 weeks. In contrast, the corresponding experi-mental substances were only administered in the last four weeks.

The experimental substances I-ol, I-an and UDCA were dissolved in a 0.8 % aqueous HPMC gel to obtain the corresponding concentrations: **(1)** I-ol with 0.4, 4 and 40 mg/kg b.w. **(2)** I-an with 0.4, 4 and 40 mg/kg b.w. **(3)** UDCA at 40 mg/kg b.w. Group 2 and 3 rats received only HPMC as placebo. Diets were fed ad libitum and their intake per cage was controlled daily. The experimantal substances or only HPMC were administered intragastrically by oral gavage. After 16 weeks of treatment the rats were anaesthetized with an intraperitoneal injection of a 5 % pentobarbital solution and then killed by aorta scarification.

#### (B) Acute toxicity study

The three ω-imidazolyl-alkyl derivatives I-ol, I-an and I-phosphocholine were suspended in 0.8% aqueous hydroxypropyl-methylcellulose gel to the following six dose levels: 10, 50, 100, 250, 500 and 1000 mg/kg b.w.. They were administered to the animals once by an oral gavage. intragastrically. The administration volume was 20 ml/kg b.w.. For further information, look at the supplement please.

### Liver histopathology

Liver samples were randomly selected, fixed in Bouin`s solution and embedded in paraffin wax. Histological sections were prepared and stained with haematoxylin and eosin (H&E staining). Further liver samples were fixed in a solution containing formaldehyde, paraformaldehyde and glutaraldehyde with subsequent post-fixation in 1 % osmium containing phosphate buffer pH 7.4 for further analysis. Liver tissue was embedded in a mixture of butyl-methyl-metacrylates. Sections of 0.5 – 1.0 µm thickness were stained with Asur II, methylene blue and alkaline fuxine.

## Results

### ROS state

I-ol reduced the HFD effect on the amount of **superoxide radical anion (SRA)** [Figure 1 (a)] by an average of 60 %, its highest concentration reducing it to 24 % below the reference value measured in healthy animals of control group 1. I-an in the middle and highest concentration reduced the amount by 54 % and 75 %. I-ol showed a strong, extremely significant linear dose-dependent effect respectively, while I-an revealed a weak, significant linear dose-dependent effect. UDCA was also able to reduce the SRA amount by 84 %.

**Figure 1.**
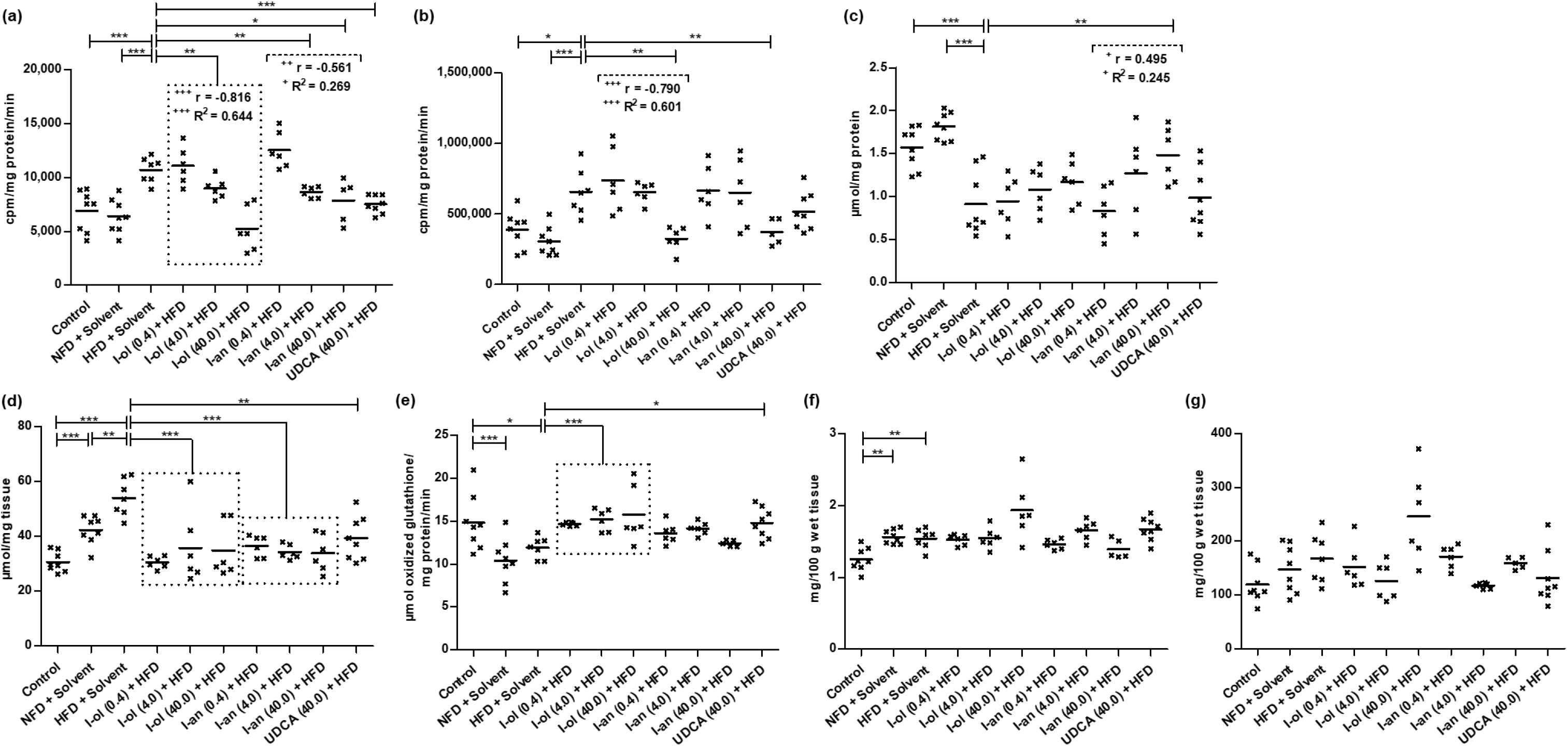
In vivo ROS state in rat liver: **(a)** SRA in HFD-fed rats is elevated by 55 % and 67 % respectively compared to control groups 1 and 2. Compared to the disease group, I-ol and I-an cause an average decrease of 21 % and 23 %. UDCA displays a decrease by 30 % **(b)** Hydrogen peroxide in HFD-fed rats is elevated by 69 % and 115 % compared to control groups 1 and 2. Compared to the disease group, I-ol and I-an cause a decrease of 50 % and 43 % respectively in the highest concentration. **(c)** GSH is reduced by 42 % and 50 % in HFD-fed rats compared to control groups 1 and 2. Compared to the disease group, I-an elevates the amount of GSH by 63 %. **(d)** TBARS is 76 % and 28 % higher in HFD-fed rats compared to control groups 1 and 2, which have a 38 % higher concentration than control group 1. I-ol and I-an show an average decrease of 38 % and 35 % respectively compared to the disease group **(e)** GR activity is decreased by 20 % in HFD-fed rats compared to control group 1, while it is increased by 14 % compared to control group 2, which shows a decrease of 30 % compared to control group 1. Compared to the disease group, I-ol increases enzymatic activity by 28 %. UDCA elevates activity by 24 %. **(f)** GPx activity in HFD-fed rats is increased by 22 % compared to control group 1, but without difference to control group 2, which has 24 % higher activity than control group 1. **(g)** Catalase activity shows no difference between all groups. Rats from the disease group (n = 8) as well as from the treatment groups (n = 6 to 8) could not avoid daily HFD intake. Each point represents the individual value of the pathologically relevant parameter of a single rat. The mean value is displayed as horizontal line. Statistical calculation was based on one-way ANOVA and subsequent complex planned contrast or simple contrast analysis. Bonferroni adjustment for multiple comparisons reduced the significance level to * p < 0.0167; ** p < 0.00333; *** p< 0.000333. Dose-dependent effects were specified by Pearson’s correlation coefficient r and linear regression analysis by the coefficient of determination R^2^ with ^+^ p < 0.05; ^++^ p < 0.01; ^+++^ p < 0.001.

I-ol and I-an in their highest concentration not only completely reduced the HFD effect on the amount of **hydrogen peroxide** [Figure 1 (b)], but decrease it by 16 % and 4 % respectively below the reference values measured in healthy animals of group 1. I-ol showed a strong, extremely significant linear dose-dependent effect. UDCA displayed no statistically significant changes.

I-an in its highest concentration reduced the HFD effect on the amount of **reduced glutathione (GSH)** [Figure 1 (c)] by 87 % and showed a weak, significant linear dose-dependent effect. I-ol displayed an insignificant trend towards an rising GSH level. UDCA was ineffective.

I-ol and I-an reduced the HFD effect on the amount of **thiobarbituric acid reactive substances (TBARS)** [Figure 1 (d)] by an average of 87 % and 82 % respectively. UDCA displayed a 63 % reduction of the HFD effect.

I-ol and UDCA completely abolished the HFD effect on the enzymatic activity of **glutathione reductase (GR)** [Figure 1 (e)], which increased the activity by 113 % and 99 % on average, while I-an showed no effect.

None of the experimental compounds showed a statistically relevant effect on the enzymatic activity of **glutathione peroxidase (GPx)** [Figure 1 (f)] or **catalase** [Figure 1 (g)].

### Lipid balance

I-ol and I-an reduced the HFD effect on the concentration of **hepatic triglycerides** [Figure 2 (a)] by an average of 41 % and 33 % respectively. UDCA diminished the HFD effect by 39 %. I-an reduced the HFD effect on the concentration of **hepatic phospholipids** [Figure 2 (b)] by an average of 53 %, with the highest concentration reducing it only to 3 % above the reference value measured in healthy animals of group 1. There was a strong, extremely significant linear dose-dependent effect. I-ol and UDCA did showed no therapeutic effect.

**Figure 2.**
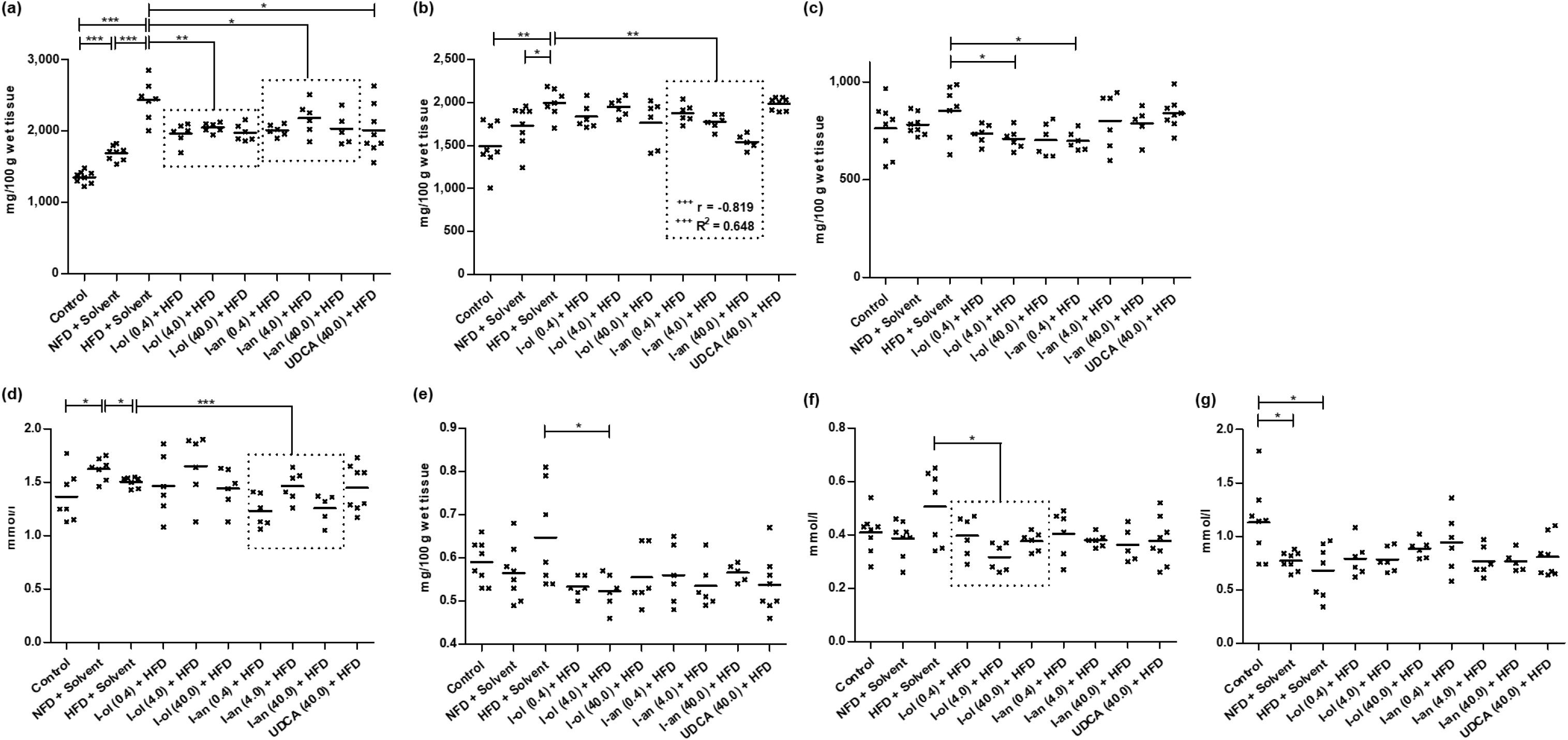
Liver and serum fat content. **(a)** Liver triglycerides in HFD-fed rats are elevated by 80 % and 44 % respectively compared to control groups 1 and 2, giving a concentration 25 % higher than control group 1. Compared to the disease group, I-ol and I-an cause an average decrease of 18 % and 15 % respectively. UDCA displays a decrease of 17 %. **(b)** Liver phospholipids are elevated by 33 % and 15 % in HFD-fed rats compared to control groups 1 and 2. Compared to the disease group, I-an causes an average decrease of 13 %. **(c)** Hepatic cholesterol shows no differences between the two control groups and the disease group. Compared to the disease group, I-ol in the middle and I-an in the low concentration reduce the amount of liver cholesterol by 17 % and 18 % respectively. **(d)** Serum triglycerides in control group 2 are elevated by 19 % compared to control group 1, while the disease group has a 7 % lower concentration than control group 2. Compared to the disease group, I-an shows an average decrease of 12 %. **(e)** Serum VLDL shows no differences between the two control groups and the disease group. I-ol in the medium concentration reduces the VLDL content by 19 %. **(f)** Serum LDL cholesterol shows no differences between the two control groups and the disease group. I-ol causes an average decrease of 28 %. **(g)** Serum HDL cholesterol in the disease group is reduced by 40 % compared to control group 1, which has a 32 % higher concentration than control group 2. Rats from the disease group (n = 8) as well as from the treatment groups (n = 6 to 8) could not avoid daily HFD intake. Each point represents the individual value of the pathologically relevant parameter of a single rat. The mean value is displayed as horizontal line. Statistical calculation was based on one-way ANOVA and subsequent complex planned contrast or simple contrast analysis. Bonferroni adjustment for multiple comparisons reduced the significance level to * p < 0.0167; ** p < 0.00333; *** p< 0.000333. Dose-dependent effects were specified by Pearson’s correlation coefficient r and linear regression analysis by the coefficient of determination R^2^ with ^+^ p < 0.05; ^++^ p < 0.01; ^+++^ p < 0.001.

Although all three concentrations of I-ol displayed a comparable reduction of the **hepatic amount of cholesterol** [Figure 2 (c)], only the middle concentration and I-an in the low concentration completely reduced the HFD effect. They even lowered the cholesterol amount by 7 % and 8 % below the reference value measured in healthy animals of group 1.

I-an not only completely reduced the HFD effect on the concentration of **serum triglycerides** [Figure 2 (d)], but even reduced it by an average of 4 % below the reference value measured in healthy animals of group 1. I-ol and UDCA showed no therapeutic influence.

Although all three experimental substances exhibited a trend towards a decreasing concentration of **serum VLDL** [Figure 2 (e)], only I-ol could not only statistically reduce the HFD effect completely, but also decreased the VLDL concentration far below the reference value measured in healthy animals of group 1.

I-ol not only completely reduced the HFD effect on the concentration of **LDL cholesterol** in serum [Figure 2 (f)], but also lowered it by an average of 11 % below the reference value measured in healthy animals of group 1. I-an and UDCA showed a statistically non-significant trend towards lowering LDL serum concentration.

None of the experimental compounds showed any statistically relevant effect on the concentration of **serum HDL cholesterol** [Figure 2 (g)].

In summary, I-ol and I-an showed a therapeutic effect in nearly all measured parameters, whereas UDCA only had a corresponding effect on the hepatic triglyceride concentration. A concentration dependence has been demonstrated for I-an in the reduction of hepatic phospholipid levels. The NFD group showed an increase in the concentration of liver and serum triglyceride and a decrease in the HDL cholesterol concentration. None of the three compounds had a therapeutic effect on HDL cholesterol concentration. There was no statistically relevant difference between the disease and both control croups regarding the concentration of liver cholesterol, serum VLDL and LDL cholesterol in serum.

### Adipocytokines (adiponectin, leptin and TNF-α) and glucose metabolism

I-ol and I-an increased the HFD effect on the concentration of **serum adiponectin** [<u>Rats from</u> the disease group (n = 8) as well as from the treatment groups (n = 6 to 8) could not avoid daily HFD intake. Each point represents the individual value of the pathologically relevant parameter of a single rat. The mean value is displayed as horizontal line. Statistical calculation was based on one-way ANOVA and subsequent complex planned contrast or simple contrast analysis. Bonferroni adjustment for multiple comparisons reduced the significance level to * p < 0.0167; ** p < 0.00333; *** p< 0.000333. Dose-dependent effects were specified by Pearson’s correlation coefficient r and linear regression analysis by the coefficient of determination R^2^ with ^+^ p < 0.05; ^++^ p < 0.01; ^+++^ p < 0.001.

Figure 3 (a)] by an average of 78 % and 38 %, respectively, with I-ol at its highest concentration increasing it to 4 % above the reference value measured in healthy animals of control group 1. UDCA elevated the concentration by 52 %. I-an showed a weak, significant linear dose-dependent effect.

**Figure 3.**
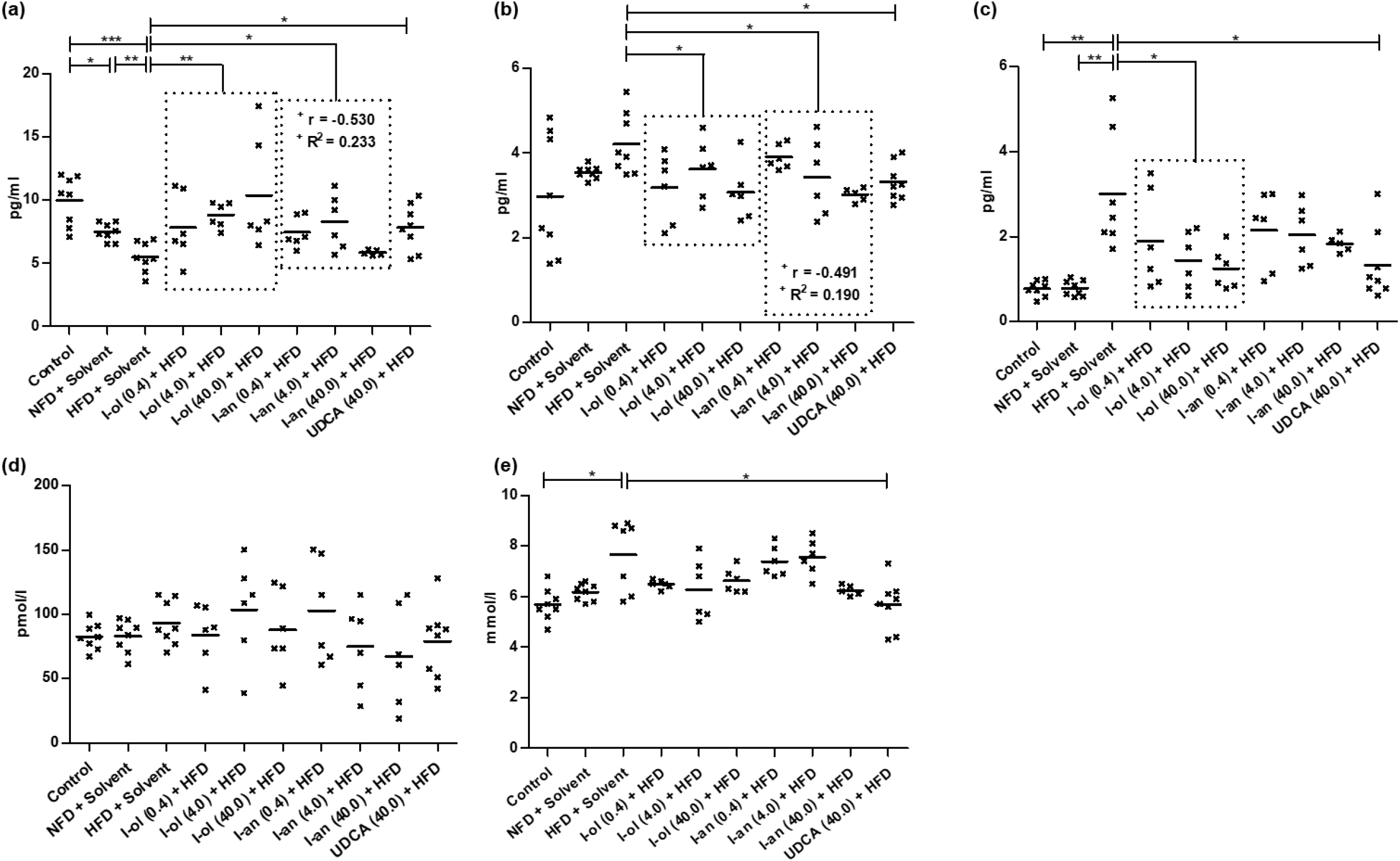
Adipocytokines and glucose metabolism: **(a)** Serum adiponectin is reduced by 45 % and 27 % respectively in HFD-fed rats compared to control groups 1 and 2, which have a concentration 25 % lower than that of control group 1. I-ol and I-an cause an average increase of 64 % and 31 % respectively compared to the disease group. UDCA shows an increase of 42 %. **(b)** Serum leptin is non-significantly elevated by 41 % and 19 % compared to control groups 1 and 2. Compared to the disease group, I-ol and I-an cause an average decrease of 22 % and 18 % respectively. UDCA shows a decline of 21 %. **(c)** Serum TNF-α is elevated by 286 % and 280 % respectively in HFD-fed rats compared to control groups 1 and 2. Compared to the disease group, I-ol causes an average decrease of 49 %. UDCA displays a decrease of 56 %. **(d)** Serum insulin concentration shows no difference between the groups. **(e)** Serum glucose in HFD-fed rats is elevated by 35 % compared to control group 1. UDCA decreases by 26%. Rats from the disease group (n = 8) as well as from the treatment groups (n = 6 to 8) could not avoid daily HFD intake. Each point represents the individual value of the pathologically relevant parameter of a single rat. The mean value is displayed as horizontal line. Statistical calculation was based on one-way ANOVA and subsequent complex planned contrast or simple contrast analysis. Bonferroni adjustment for multiple comparisons reduced the significance level to * p < 0.0167; ** p < 0.00333; *** p< 0.000333. Dose-dependent effects were specified by Pearson’s correlation coefficient r and linear regression analysis by the coefficient of determination R^2^ with ^+^ p < 0.05; ^++^ p < 0.01; ^+++^ p < 0.001.

I-ol and I-an decreased the HFD effect on the concentration of **serum leptin** [<u>Rats from</u> the disease group (n = 8) as well as from the treatment groups (n = 6 to 8) could not avoid daily HFD intake. Each point represents the individual value of the pathologically relevant parameter of a single rat. The mean value is displayed as horizontal line. Statistical calculation was based on one-way ANOVA and subsequent complex planned contrast or simple contrast analysis. Bonferroni adjustment for multiple comparisons reduced the significance level to * p < 0.0167; ** p < 0.00333; *** p< 0.000333. Dose-dependent effects were specified by Pearson’s correlation coefficient r and linear regression analysis by the coefficient of determination R^2^ with ^+^ p < 0.05; ^++^ p < 0.01; ^+++^ p < 0.001.

Figure 3 (b)] by 75 % and 62 % on average, with highest concentration reducing it to 3 % and 1 % above the reference value measured in healthy animals of control group 1. UDCA reduced the concentration by 72 %. I-an showed a weak, significant linear dose-dependent effect.

I-ol reduced the HFD effect on the concentration of **serum TNF-α** [<u>Rats from</u> the disease group (n = 8) as well as from the treatment groups (n = 6 to 8) could not avoid daily HFD intake. Each point represents the individual value of the pathologically relevant parameter of a single rat. The mean value is displayed as horizontal line. Statistical calculation was based on one-way ANOVA and subsequent complex planned contrast or simple contrast analysis. Bonferroni adjustment for multiple comparisons reduced the significance level to * p < 0.0167; ** p < 0.00333; *** p< 0.000333. Dose-dependent effects were specified by Pearson’s correlation coefficient r and linear regression analysis by the coefficient of determination R^2^ with ^+^ p < 0.05; ^++^ p < 0.01; ^+++^ p < 0.001.

Figure 3 (c)] by an average of 66 % and UDCA by 75 %. I-an displayed an insignificant trend towards a falling TNF-α level.

While the concentration of **serum insulin** [<u>Rats from</u> the disease group (n = 8) as well as from the treatment groups (n = 6 to 8) could not avoid daily HFD intake. Each point represents the individual value of the pathologically relevant parameter of a single rat. The mean value is displayed as horizontal line. Statistical calculation was based on one-way ANOVA and subsequent complex planned contrast or simple contrast analysis. Bonferroni adjustment for multiple comparisons reduced the significance level to * p < 0.0167; ** p < 0.00333; *** p< 0.000333. Dose-dependent effects were specified by Pearson’s correlation coefficient r and linear regression analysis by the coefficient of determination R^2^ with ^+^ p < 0.05; ^++^ p < 0.01; ^+++^ p < 0.001.

Figure 3 (d)] showed no difference between all groups, UDCA completely reduced the HFD effect on the concentration of **serum glucose** [<u>Rats from</u> the disease group (n = 8) as well as from the treatment groups (n = 6 to 8) could not avoid daily HFD intake. Each point represents the individual value of the pathologically relevant parameter of a single rat. The mean value is displayed as horizontal line. Statistical calculation was based on one-way ANOVA and subsequent complex planned contrast or simple contrast analysis. Bonferroni adjustment for multiple comparisons reduced the significance level to * p < 0.0167; ** p < 0.00333; *** p< 0.000333. Dose-dependent effects were specified by Pearson’s correlation coefficient r and linear regression analysis by the coefficient of determination R^2^ with ^+^ p < 0.05; ^++^ p < 0.01; ^+++^ p < 0.001.

Figure 3 (e)] by 100 %. I-ol and I-an had no therapeutical effect.

### Liver enzymes, cholestase and fibrosis parameters

In this study, the activity of ALT and AST increased by 41 % and 30 %, respectively, over control group 1, while the De-Ritis quotient was 1.8 indicating severe liver damage in the disease group.

I-ol reduced the HFD effect on the enzymatic activity of **ALT** [Figure 4 (a)] by an average of 55 %, in the highest concentration even by 66 %. I-an reduced this effect in its highest concentration by 68 % and showed a week, significant linear dose-dependent effect. UDCA displayed a 50 % reduction in the HFD effect.

**Figure 4.**
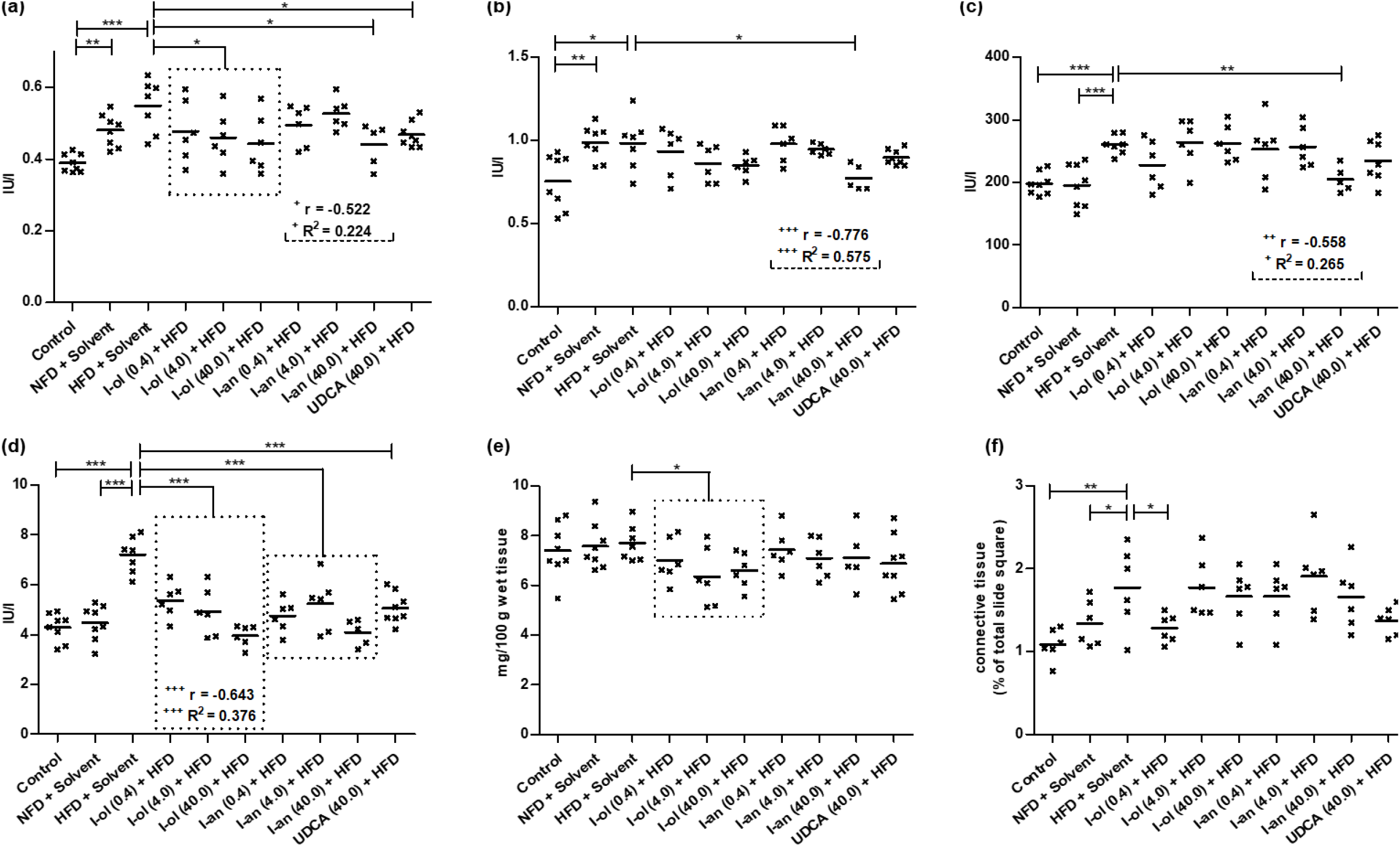
Liver enzymes, cholestase and fibrosis parameters: **(a)** ALT in HFD-fed rats is elevated by 41 % compared to control group 1, which has 23 % lower enzymatic activity compared to control group 2. Compared to the disease group, I-ol shows an average decrease of enzymatic activity by 16 % and I-an in the highest concentration by 20%. UDCA shows of reduction by 15 %. **(b)** AST is elevated by 30 % in HFD-fed rats compared to control group 1, which has 31 % less enzymatic activity compared to control group 2. Compared to the disease group, I-an shows a 21 % decrease in enzymatic activity. **(c)** AP in HFD-fed rats is increased by 32 % and 33 % compared to control groups 1 and 2. Compared to the disease group, I-an at its highest concentration shows an average 21 % decrease of enzymatic activity. **(d)** γ-GT is elevated by 68 % and 61 % in HFD-fed rats compared to control groups 1 and 2. Compared to the disease group, I-ol and I-an show an average decrease in enzymatic activity by 34 % and 35 % respectively. UDCA shows a reduction of 30 %. **(e)** Total serum bilirubin shows no differences between the two control groups and the disease group. Compared to the disease group, I-ol shows an average decrease in concentration of 14 %. **(f)** The square of connective tissue in the liver is increased by 63 % and 32 % respectively in HFD-fed rats compared to control groups 1 and 2. Compared to the disease group, I-ol shows a decrease by 28 % in the lowest concentration. Rats from the disease group (n = 8) as well as from the treatment groups (n = 6 to 8) could not avoid daily HFD intake. Each point represents the individual value of the pathologically relevant parameter of a single rat. The mean value is displayed as horizontal line. Statistical calculation was based on one-way ANOVA and subsequent complex planned contrast or simple contrast analysis. Bonferroni adjustment for multiple comparisons reduced the significance level to * p < 0.0167; ** p < 0.00333; *** p< 0.000333. Dose-dependent effects were specified by Pearson’s correlation coefficient r and linear regression analysis by the coefficient of determination R^2^ with ^+^ p < 0.05; ^++^ p < 0.01; ^+++^ p < 0.001.

I-an in its highest concentration prevented the HFD effect on the enzymatic activity of **AST** [Figure 4 (b)] <u>almost completely</u> by a 92 % reduction and showed a moderate, extremely significant linear dose-dependent effect. I-ol and UDCA showed no therapeutic influence.

I-an int its highest concentration reduced the HFD effect on the enzymatic activity of **alkaline phosphatase (AP)** [Figure 4 (c)] by 89 %. It displayed a weak, significant linear dose-dependent effect. I-ol and UDCA showed no therapeutic influence.

I-ol and I-an reduced the HFD effect on the enzymatic activity of **γ-glutamyltransferase (γ-GT)** [Figure 4 (d)] by 84 % and 86 %, respectively, which corresponded to an almost complete elimination. I-ol showed a moderate, extremely significant linear dose-dependent effect. UDCA displayed a 73 % reduction of the HFD effect.

I-ol reduced the HFD effect on the concentration of **total bilirubin** [Figure 4 (e)] on average by more than 10 **%** below the disease and the two control groups. I-an and UDCA showed no therapeutic influence.

I-ol at in lowest concentration reduced the HFD effect on the **liver square of connective tissue** [Figure 4 (f)] by 72 %. I-an and UDCA showed no therapeutic effect.

In summary, the generally extremely significant increase of all four liver enzyme activities could be nearly completely reduced by all tested compounds in a usually very or extremely significant way. There was a trend towards a maximum effect at the lowest concentration of I-ol and I-an, which showed a weak or moderate linear dose-dependent effect.

### Histopathology

Rats fed HFD developed macrovesicular steatosis [Figure 5, Supplement Figure 2 b + c] with focal necrosis [Supplement Figure 2 c] and intralobular lymphocytic infiltration. Both ROS inhibitors (40 mg/kg b.w.) improved liver histology in NASH animals decreasing steatosis and in-flammation. UDCA seemed to have to opposite effect [Figure 5 g].

**Figure 5.**
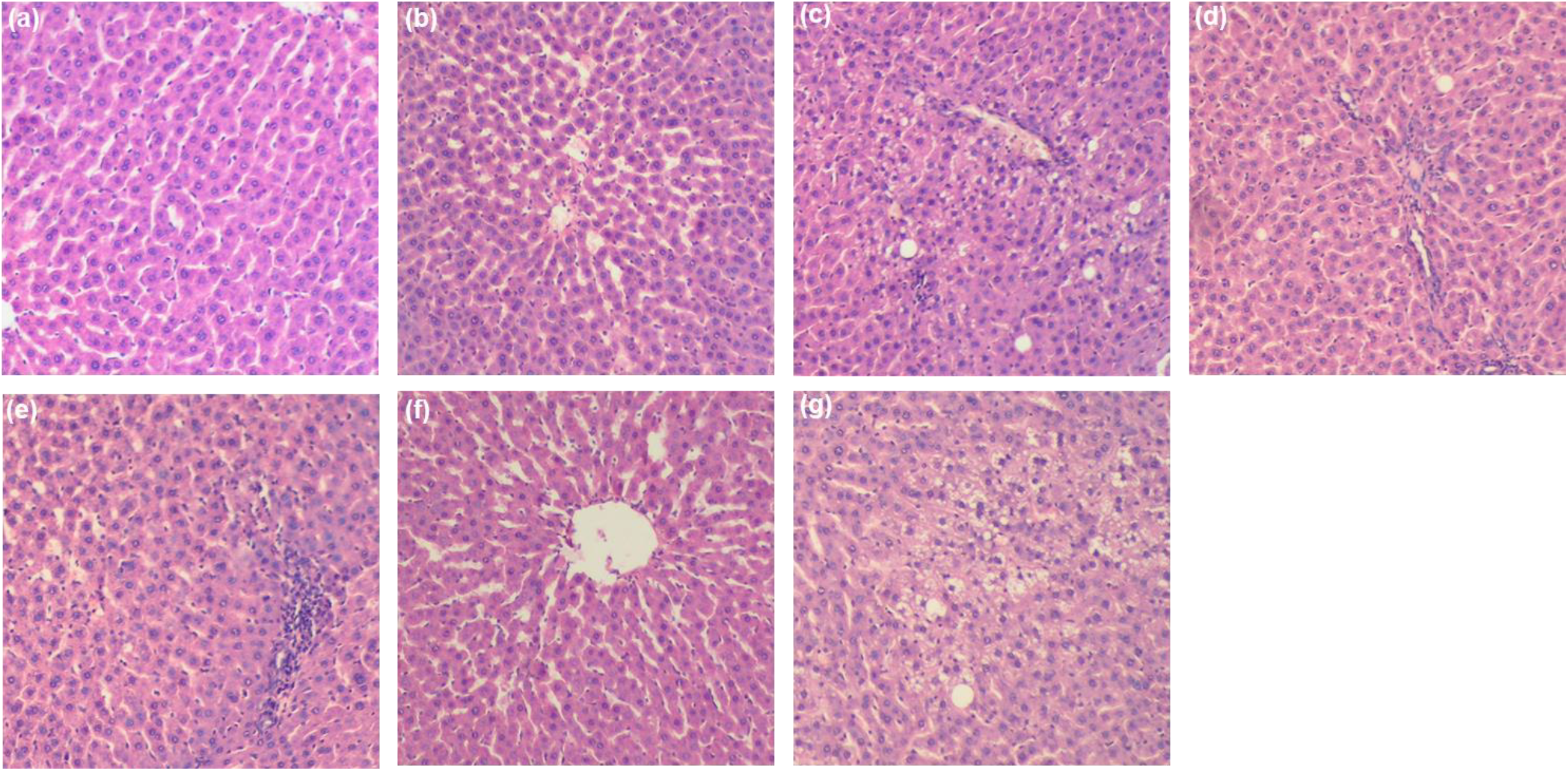
Representative histological pictures of liver sections: **(a)** Control. **(b)** NFD + Solvent. **(c)** HFD + Solvent. **(d)** I-ol (0.4 mg/kg b.w.) + HFD. **(e)** I-ol (4 mg/kg b.w.) + HFD. **(f)** I-ol (40 mg/kg b.w.) + HFD. **(g)** UDCA (40 mg/kg b.w.) + HFD. For reasons of redundancy the illustrations of the I-an groups were omitted. Nine sections of each liver were prepared – with six or eight animals per group. The H&E stained sections were shown in 10-fold magnification.

### Acute toxicity study

I-ol [Supplement Table 2] showed no acute toxicity sign up to a concentration of 1000 mg/kg b. w.. From this maximum dosage a reduced motility was observed with reduced muscle tone, ataxia and dyspnea, whereas no inhibition of body weight gain or necropsy findings were detected further on. None of the 30 rats died, so the lowest lethal dose as well as the LD_50_ dose (14 days) had to be greater than 1000 mg/kg b.w.

The application of I-an [Supplement Table 3] led to a premature death of three rates: Two animals died at a concentration of 500 mg/kg b.w. after six days and one animal at 1000 mg/kg b.w. after five days. Thus, the lowest lethal dose was 500 mg/kg b.w., whereas the LD_50_ dose (14 days) had to be greater than 1000 mg/kg b.w..

I-phosphocholine [Supplement Table 4] showed no signs of acute toxicity up to a concentration of 100 mg/kg b. w.. At 250 mg/kg b.w. slight to moderate reduced motility, ataxia and dyspnea were recorded in all five rats within a period of 30 minutes to three days. Within a period of three hours to two days, slightly reduced muscle tone and impaired piloerection were additional toxic signs in all five rats receiving a dose of 500 mg/kg b. w.. The increase of the dosage to 1000 mg/kg b.w. led to a significant reduction of muscle tone and additionally to ptosis and chromodacryorrhea after two days. One rat died after two days and three more after three days. Thus, the lowest lethal dose was 1000 mg/kg b.w., whereas the LD_50_ dose (14 days) was 880 mg/kg b.w..

A summary of the study results is depicted in [Table 1].

**Table 1.**
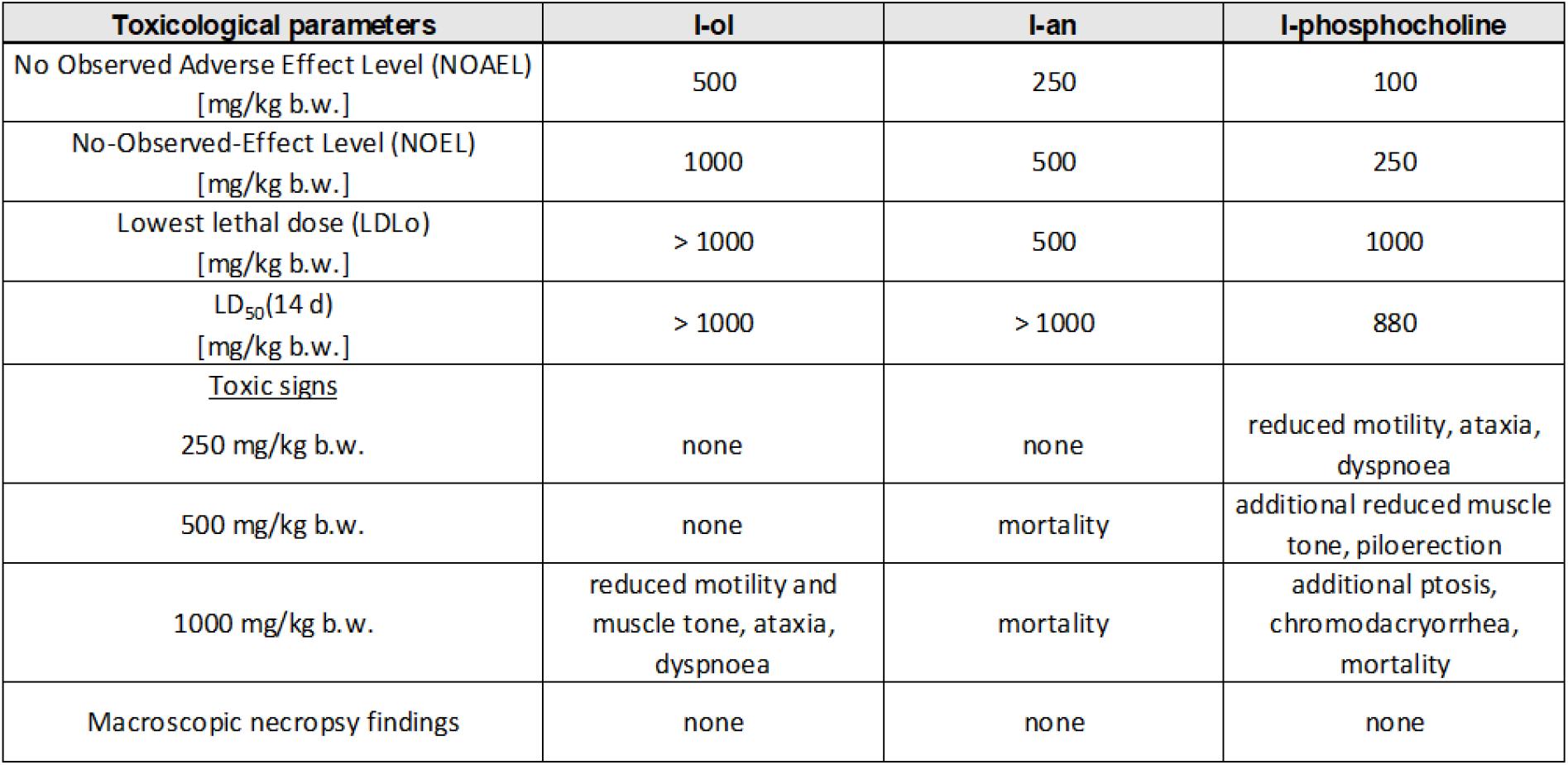
Summary of the acute toxicity study with the ω-imidazolyl-alkyl derivatives

| Toxicological parameters | I-ol | I-an | I-phosphocholine |
| --- | --- | --- | --- |
| No Observed Adverse Effect Level (NOAEL)<br>[mg/kg b.w.] | 500 | 250 | 100 |
| No-Observed-Effect Level (NOEL)<br>[mg/kg b.w.] | 1000 | 500 | 250 |
| Lowest lethal dose (LDLo)<br>[mg/kg b.w.] | > 1000 | 500 | 1000 |
| LD<sub>50</sub>(14 d)<br>[mg/kg b.w.] | > 1000 | > 1000 | 880 |
| <u>Toxic signs</u><br>250 mg/kg b.w. | none | none | reduced motility, ataxia,<br>dyspnoea |
| 500 mg/kg b.w. | none | mortality | additional reduced muscle<br>tone, piloerection |
| 1000 mg/kg b.w. | reduced motility and<br>muscle tone, ataxia,<br>dyspnoea | mortality | additional ptosis,<br>chromodacryorrhea,<br>mortality |
| Macroscopic necropsy findings | none | none | none |

## Discussion

We have already shown in an animal model that the new CYP2E1 enzyme inhibitors I-ol and I-an are excellent drug candidates for the treatment of alcoholic steatohepatitis. Here we wanted to investigate whether they are just as suitable for a therapy of NASH.

The enzymatic activities of ALT and AST in serum are described as elevated in patients with steatosis and NASH. However, this observation is not mandatory(16). The diagnostic value of ALT for NASH is still controversial and is based on unclear study results(17). In our study, the activity of ALT in rats in the HFD group is increased compared to rats in the NFD group. This observation is supported by the results of another study, in which the activity of ALT reached its maximum in week 16, while the development of steatohepatitis began in week 12(18). The increase of ALT was diminished by almost 70 % by I-ol and I-an, while UDCA had a lesser effect. I-an even achieved a 90 % reduction in AST. Studies with NASH patients showed only an insignificant reduction of ALT by UDCA(19).

Increased alkaline phosphatase (AP) levels observed in the HFD group were also reported several times for patients with NASH(17). Only I-an reduced the enhanced enzyme activity, but this almost completely.

The serum level of γ glutamyltransferase (GGT) in animals in the HFD group was extremely significantly elevated. Basically, GGT is described as an early predictive marker for insulin resistance, NAFLD, metabolic syndrome and many other human diseases(20). In particular, GGT is thought to play an important role in the defense mechanism against oxidative stress by being involved in the extracellular transport and cellular uptake of GSH(21). Thus it can be concluded that the animals of the HFD group were exposed to massive oxidative stress.

Paying attention to the fact that NASH can also develop fibrosis, we determined the amount of connective tissue in liver sections. The liver of HFD-fed animals showed an increase in connective tissue compared to both control groups, suggesting an induction of fibrosis. This observation has only been described in the literature after week 24(18). Only I-ol delivered a first clue of a possible antifibrotic effect of the drug candidates.

Adiponectin and TNF-α should play a significant role in the development of NASH(22). Rats of the HFD group showed reduced adiponectin levels associated with insulin resistance(23) as well as increased serum levels of TNF-α pointing to NASH. These two tissue hormones and leptin influence each other in an antagonistic way.

The concentration of adiponectin is decreased in the serum of rats from the HFD group, albeit they have not gained body weight, or their insulin and glucose levels have not or hardly been increased. Thus, there were no signs of advanced overweight or manifestation of insulin resistance or even type 2 diabetes in these animals. In the case of ASH, it was reported that a reduced adiponectin concentration is primarily related to oxidative stress caused by CYP2E1(24). In addition, the effect of alcohol on fatty tissue is well documented. However, there are few reports on the association of NASH and the importance of CYP2E1 in adipose tissue. A study on a leptin deficient mouse model for diabetes mellitus concluded that the concentration of adiponectin already decreased at the transition of non-alcoholic fatty liver to NASH, whereas it remained constant without the presence of steatosis(25). However, the degree of steatosis correlated with the body weight of the animals. I-ol restored the adiponectin concentration in serum to the level of rats from the control group in a concentration-dependent manner. I-an and UDCA had a slightly smaller effect.

A higher level of TNF-α in patients with NAFLD and NASH has been described in numerous publications, but this marker does not allow staging of NAFLD(17). In particular, I-ol reduced the elevated TNF-α level dose-dependently and thus one of the prominent surrogate markers for NAFLD.

The serum concentration of leptin in HFD-fed rats should be permanently increased from week 1 - 16 compared to the standard diet(26). However, there is currently a disagreement as to whether this is also the case for patients with NASH(22). Our study showed a strong but statistically not significant increase in leptin concentration in rats from the disease group compared to those from the control group. I-ol and I-an reduced this HFD effect in their highest concentration almost to the initial level of the control group.

All animals of the HFD group showed only a partial change of the lipid parameters of liver and serum compared to those of the control group. In particular, the amount of triglycerides in the liver was significantly increased in accordance with study results in NAFLD patients(27). This effect was reduced by our drug candidates. Only I-an reduced below the levels of both reference groups, whereas I-ol and UDCA showed no effect. The situation was similar to the remaining lipid parameters such as hepatic cholesterol and phospholipids, VLDL as well as LDL und HDL cholesterol in serum, where no clear statement could be made regarding both the disease severity of the animals in the HFD group and the therapeutic success of the drug candidates. This finding is supported by the lack of increase in body and liver weight in the disease group (data not shown). However, these partially contradictory results reflect the current state of scientific literature. There was no increase in triglycerides in serum^(18)^ and no gain in body weight(28) in the HFD group compared to the control group. This observation was contradicted by results from other animal experiments in which solid food was used and an increase in liver weight was recorded weeks before an increase in body weight(18). In our study, the lack of increase in body weight may be due to the basic components of the liquid diet. HPMC or guar gum led to reduced weight gain in rats as part of a high-fat diet(29). Comparable results have been described with other species. The results of our histopathological examinations of the liver showed, in agreement with the observations from other experiments(30) but modest compared to human patients in a final stage(31), a pronounced macrovesicular steatosis, massive necrotic areas and infiltrations by macrophages and lymphocytes in the disease group. I-ol in particular led to a dose-dependent reduction of pathological signs. First steatotic vesicles disappeared with increasing concentration of I-ol then finally infiltration with mononuclear cells. UDCA showed no effect.

In contrast to other study results(27), we found that the amount of cholesterol in the liver of animals of the HFD group was only slightly and statistically not significantly increased compared to the control groups. In another study in histologically confirmed NASH patients, a correlation was established between the severity of NAFLD and the amount of hepatic cholesterol(32). The driving force was assumed to be the dysregulation of HMG-CoA reductase and the associated accumulation of free cholesterol, which was not measured in our study. The discrepancy in these results could be due to the export of hepatic cholesterol from the liver. Furthermore, disease progression in the HFD group may not have reached the stage where cholesterol accumulates in the liver, or our animal model may not be suitable in this respect to compare its results with studies in NASH patients. Further explanations are the cholesterol-free nutrition in our animal model and the composition of the liquid nutrition. UDCA increases the amount of hepatic cholesterol through increased synthesis in patients with pathological obesity(33). This observation could explain the lack of an effect of UDCA in our study, whereas our compounds were able to reduce the amount of hepatic cholesterol below the control groups.

In our study the LDL concentration in the serum of the disease group was only slightly and statistically not significantly increased compared to the control groups, whereas the HDL concentration showed a statistically significant decrease. A decrease in HDL levels has been observed in obese patients with NAFLD(32) and, in principle, in patients with NASH(34). I-ol was able to significantly lower LDL levels, whereas neither our drug candidates nor UDCA had any effect on HDL levels. VLDL concentration in serum is significantly higher in patients with steatosis or NASH(34). Animals in our disease group have non-significantly elevated serum levels of VLDL, with I-ol, I-an and UDCA appearing to have a reducing effect on the release of VLDL by the liver.

All measured parameters of ROS stress in the livers of rats of the disease group were changed at least very significantly compared to the control groups - with the exception of the enzymatic activity of catalase. In particular, this concerned superoxide radical anion, hydrogen peroxide, equivalents of lipid peroxidation and reduced glutathione. Especially for reduced glutathione, the reduction was so severe that an exhaustion of the cellular protective system against ROS had to be assumed. Thus, it could be assumed that the feeding with HFD caused a severe burden of the liver with ROS stress. Both drug candidates were able to relieve ROS stress in a very significant and partly dose-dependently way. UDCA basically had a weaker effect and no influence on the amount of hydrogen peroxide and reduced GSH. This observation was consistent with study results that denied UDCA any ROS-protective effect in patients with NAFLD(35).

The importance of CYP2E1 as an essential pathological source of intracellular ROS is widely recognized in literature. The animal model we used showed a significant increase in mRNA and protein levels of CYP2E1 in rat liver(28)^,(36)^. Increased enzymatic activity of CYP2E1 in patients with NASH has also been described(37)^,(38),(39),(40)^., whereas no difference in mRNA levels between patients with steatosis and NASH could be observed(37). We have already shown in in-vitro experiments that I-ol and I-an are strong inhibitors of CYP2E1. Overall, these facts suggest that almost all of the ROS stress that arises in the high-fat diet in our experiments is due to CYP2E1.

In the acute toxicity study with rats, which was conducted according to ICH guidelines, no acute effect up to a dosage of 500 g/kg b.w. was observed for both drug candidates. However, two of the five rats that got I-an died. I-ol did not lead to any death. Another ω-imidazolyl-alkyl derivative was tested as part of the rational drug development process: I-phosphocholine, which already showed first signs of acute toxicity at 250 g/kg b.w.. The LD_50_ dosage was estimated to be beyond 1000 mg/kg b.w. for both experimental substances, whereas 880 mg/kg b.w. was calculated for I-phosphocholine. These observations could be correlated with the binding strength of these inhibitors to CYP2E1, according to which I-ol had the highest affinity, followed by I-an and the significantly weaker acting I-phosphocholine. This could explain the different acute signs of toxicity in the sense of off-target effects. Overall, the acute toxic effect of all three substances was so low that no classification according to the usual criteria could be used(41).

In summary, the pathological parameters an histological staining used in this study indicate that the Wistar rats of our HFD disease group developed a non-alcoholic fatty liver disease (NAFLD) in general and NASH in particular, despite the fact that the surrogate markers cannot distinguish between steatosis and NASH. Our two inhibitors influenced the degree of NASH, partly dose-dependently, by attenuating the associated disease parameters. This particularly affected the ROS status of the liver, which was mainly caused by the increased enzymatic activity of CYP2E1 in the course of disease. Both CYP2E1 inhibitors could be classified as non-acute toxic. In contrast to other potential drug targets, they intervene at a suitable molecular site in the development path of NAFLD, in particular NASH They are therefore promising candidates for drug treatment of NASH, but further research is needed.

## AUTHOR CONTRIBUTIONS

T. D., A. L & V. B. designed and performed the experiments, analyzed the data, supervised the project and wrote the manuscript. O. L. provided the in vivo experiments and wrote the manuscript. E. B. and V. A. provided histological examination. E. N. was responsible for the analysis of lipid related parameters as well as establishment of the animal model. R. D., S. K., D. B., S. B. & C. R. provided technical support. D. M.-E. developed and synthesized the ω-imidazolyl-alkyl derivatives and designed the experiments. T. W. supervised the project. T. H. founded and supervised the project. All authors have reviewed the manuscript.

---

**Supplement Figure 1.**
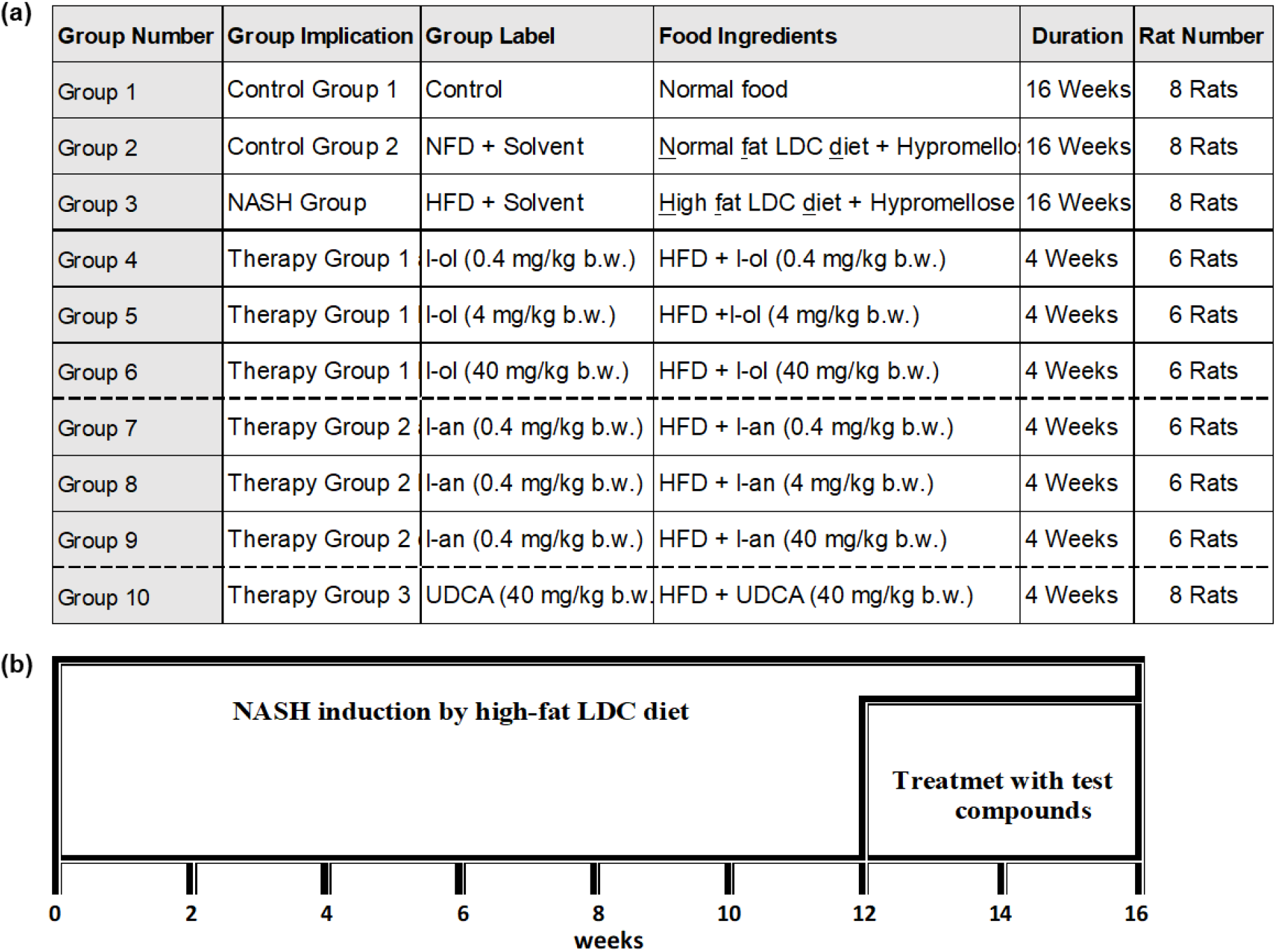
Scheme of animal experiments. **(a) Group differences:** Rats were divided into two control groups, one disease group and seven treatment groups. The disease group and the treatment groups received the HFD diet, which contained 4.9 % (m/m) corn oil. **(b) Time frame:** The complete animal study lasted 16 weeks. While all diets were administered over the entire period, the corresponding test compounds were only administered in the last four weeks. The tested compounds and Hypromellose were administered daily with oral gavage.

**Supplement Figure 2.**
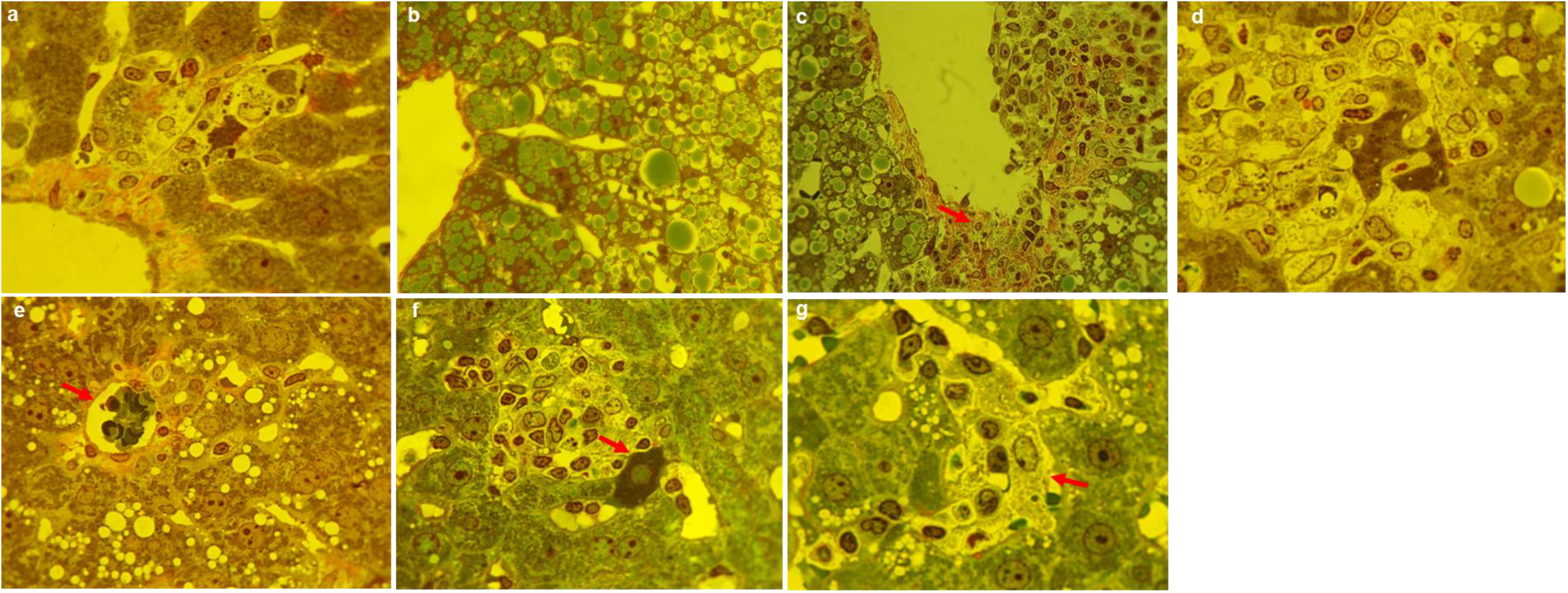
Additional histopathological pictures: Pieces of the liver were fixed in a solution containing formaldehyde, paraformaldehyde and glutaric aldehyde with a subsequent post-fixation in 1% osmium in phosphate buffer, pH 7.4. Liver tissue was embedded in a mixture of buthyl-methyl-metacrylates. Semifine sections 0.5 – 1.0 µm were stained with Asur II, methylen blue and based on fuxine. The magnification of the objective was 100 - fold and of the ocular was 10-fold **(a)** Healthy animal of the NFD group. **(b) – (d)** Animals of the HFD treated group **(b)** Hepatocytes are overloaded with manifold lipid droplets. **(c)** pronounced macrovesicular steatosis and spacious necrotic area near central vein. Nuclei of macrophages and lymphocytes are visible (arrows). **(d)** Necrotic focus. In the center are hepatocytes lysed by macrophages and lymphocytes. **(e)** Rat treated with UDCA (40 mg/kg b.w.). Inflammatory area around a hepatocyte, which is destroyed by large lipid droplet (arrow). **(f)** Rat treated with I-ol (40 mg/kg b.w.), apototic cell (arrow) on the border of a necrotic focus in portal area. **(g)** Rat treated with I-an (40 mg/kg b.w.), hypertrophic macrophage (arrow) and clustering of lymphocytes in a liver sinusoid.

**Supplement Table 1.** Composition of the liquid Lieber-De Carli (LDC) liquid diet (Lieber & De Carli, 2004)

| Compounds | LDC diet with normal fat content |  | NASH diet |  |
| --- | --- | --- | --- | --- |
|  | (g) | (%) | (g) | (%) |
| Casein | 41.4 | 4.1 | 41.4 | 4.1 |
| L-Cysteine | 0.5 | 0.1 | 0.5 | 0.1 |
| DL-Methionine | 0.3 | 0.0 | 0.3 | 0.0 |
| Maltodextrin | 25.6 | 2.6 | 25.6 | 2.6 |
| Cellulose | 10.0 | 1.0 | 10.0 | 1.0 |
| Minerals-VM | 8.8 | 0.9 | 8.8 | 0.9 |
| Vitamins-VM | 2.5 | 0.3 | 2.5 | 0.3 |
| Choline chloride* | 0.5 | 0.1 | 0.5 | 0.1 |
| Guar gum** | 3.0 | 0.3 | 3.0 | 0.3 |
| **Corn oil** | 8.5 | 0.9 | 8.5 + **40.0** | 4.9 |
| Olive oil | 28.4 | 2.8 | 28.4 | 2.8 |
| Safflower oil | 2.7 | 0.3 | 2.7 | 0.3 |
| Total | 132.2 | 13.2 | 172.2 | 17.2 |
| **Water** | **867.8** | 86.8 | **827.8** | 82.8 |
| Total | 1000.0 |  | 1000.0 |  |

\* Originally choline bitartrate, replaced by choline chloride
\*\* Originally Xanthan gum, replaced by Guar gum.

**Supplement Table 2.** Acute toxicity study results of I-ol

| Symptoms | Control | I-ol [mg/kg b.w., p.o.] |  |  |  |  |  |
| --- | --- | --- | --- | --- | --- | --- | --- |
|  | males<br>(n = 5) | 10<br>males<br>(n = 5) | 50<br>males<br>(n = 5) | 100<br>males<br>(n = 5) | 250<br>males<br>(n = 5) | 500<br>males<br>(n = 5) | 1000<br>males<br>(n = 5) |
| reduced motility | none | none | none | none | none | none | +<br>3h-6h<br>(5) |
| ataxia | none | none | none | none | none | none | +<br>3h-6h<br>(5) |
| reduced muscle tone | none | none | none | none | none | none | +<br>3h-6h<br>(5) |
| dyspnoea | none | none | none | none | none | none | +<br>3h-6h<br>(5) |
| ptosis | none | none | none | none | none | none | none |
| chromodacryorrhea | none | none | none | none | none | none | none |
| piloerection | none | none | none | none | none | +<br>2d<br>(5) | none |
| mortality |  |  |  |  |  |  |  |
| within 6 h | 0 | 0 | 0 | 0 | 0 | 0 | 0 |
| within 24 h | 0 | 0 | 0 | 0 | 0 | 0 | 0 |
| within 7 h | 0 | 0 | 0 | 0 | 0 | 0 | 0 |
| within 14 h | 0 | 0 | 0 | 0 | 0 | 0 | 0 |
| mean body weight [g] |  |  |  |  |  |  |  |
| start | 215.8 | 221.6 | 227.8 | 230.0 | 223.0 | 228.0 | 221.6 |
| after 7 d | 269.8 | 259.2 | 278.4 | 272.6 | 266.8 | 294.2 | 271.2 |
| end | 312.0 | 288.0 | 310.4 | 305.8 | 299.2 | 334.2 | 302.6 |
| mean body weight gain [%] |  |  |  |  |  |  |  |
| after 7 d | +25.0 | +17.0 | +22.2 | +18.5 | +19.6 | +29.0 | +22.4 |
| end | +44.6 | +30.0 | +36.3 | +33.0 | +34.2 | +46.6 | +36.6 |
| inhibition of body weight gain | none | none | none | none | none | none | none |
| necropsy findings | none | none | none | none | none | none | none |

**Supplement Table 3.**
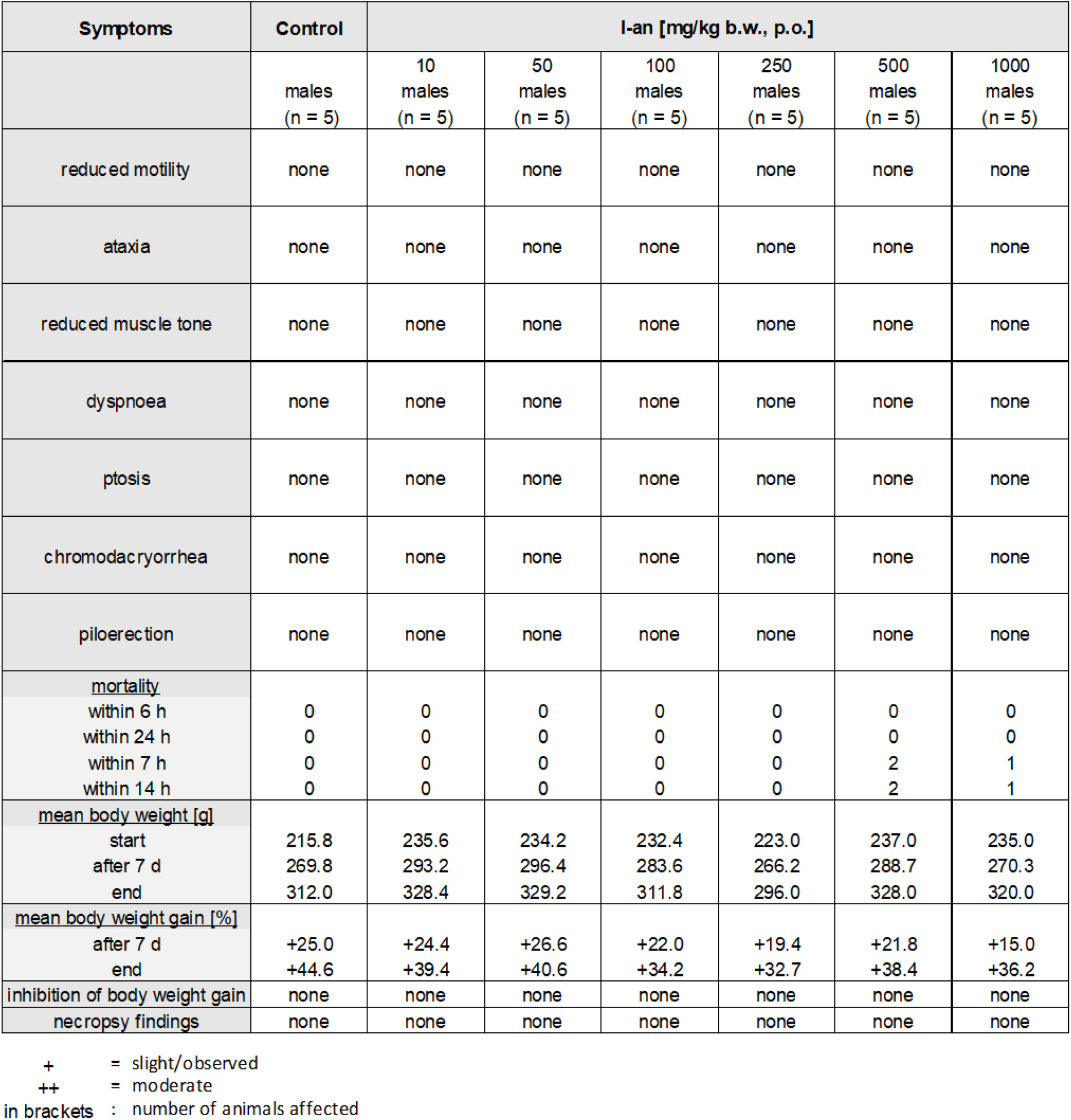
Acute toxicity study results of I-an

| Symptoms | Control | I-an [mg/kg b.w., p.o.] |  |  |  |  |  |
| --- | --- | --- | --- | --- | --- | --- | --- |
|  | males<br>(n = 5) | 10<br>males<br>(n = 5) | 50<br>males<br>(n = 5) | 100<br>males<br>(n = 5) | 250<br>males<br>(n = 5) | 500<br>males<br>(n = 5) | 1000<br>males<br>(n = 5) |
| reduced motility | none | none | none | none | none | none | none |
| ataxia | none | none | none | none | none | none | none |
| reduced muscle tone | none | none | none | none | none | none | none |
| dyspnoea | none | none | none | none | none | none | none |
| ptosis | none | none | none | none | none | none | none |
| chromodacryorrhea | none | none | none | none | none | none | none |
| piloerection | none | none | none | none | none | none | none |
| mortality |  |  |  |  |  |  |  |
| within 6 h | 0 | 0 | 0 | 0 | 0 | 0 | 0 |
| within 24 h | 0 | 0 | 0 | 0 | 0 | 0 | 0 |
| within 7 h | 0 | 0 | 0 | 0 | 0 | 2 | 1 |
| within 14 h | 0 | 0 | 0 | 0 | 0 | 2 | 1 |
| mean body weight [g] |  |  |  |  |  |  |  |
| start | 215.8 | 235.6 | 234.2 | 232.4 | 223.0 | 237.0 | 235.0 |
| after 7 d | 269.8 | 293.2 | 296.4 | 283.6 | 266.2 | 288.7 | 270.3 |
| end | 312.0 | 328.4 | 329.2 | 311.8 | 296.0 | 328.0 | 320.0 |
| mean body weight gain [%] |  |  |  |  |  |  |  |
| after 7 d | +25.0 | +24.4 | +26.6 | +22.0 | +19.4 | +21.8 | +15.0 |
| end | +44.6 | +39.4 | +40.6 | +34.2 | +32.7 | +38.4 | +36.2 |
| inhibition of body weight gain | none | none | none | none | none | none | none |
| necropsy findings | none | none | none | none | none | none | none |

**Supplement Table 4.**
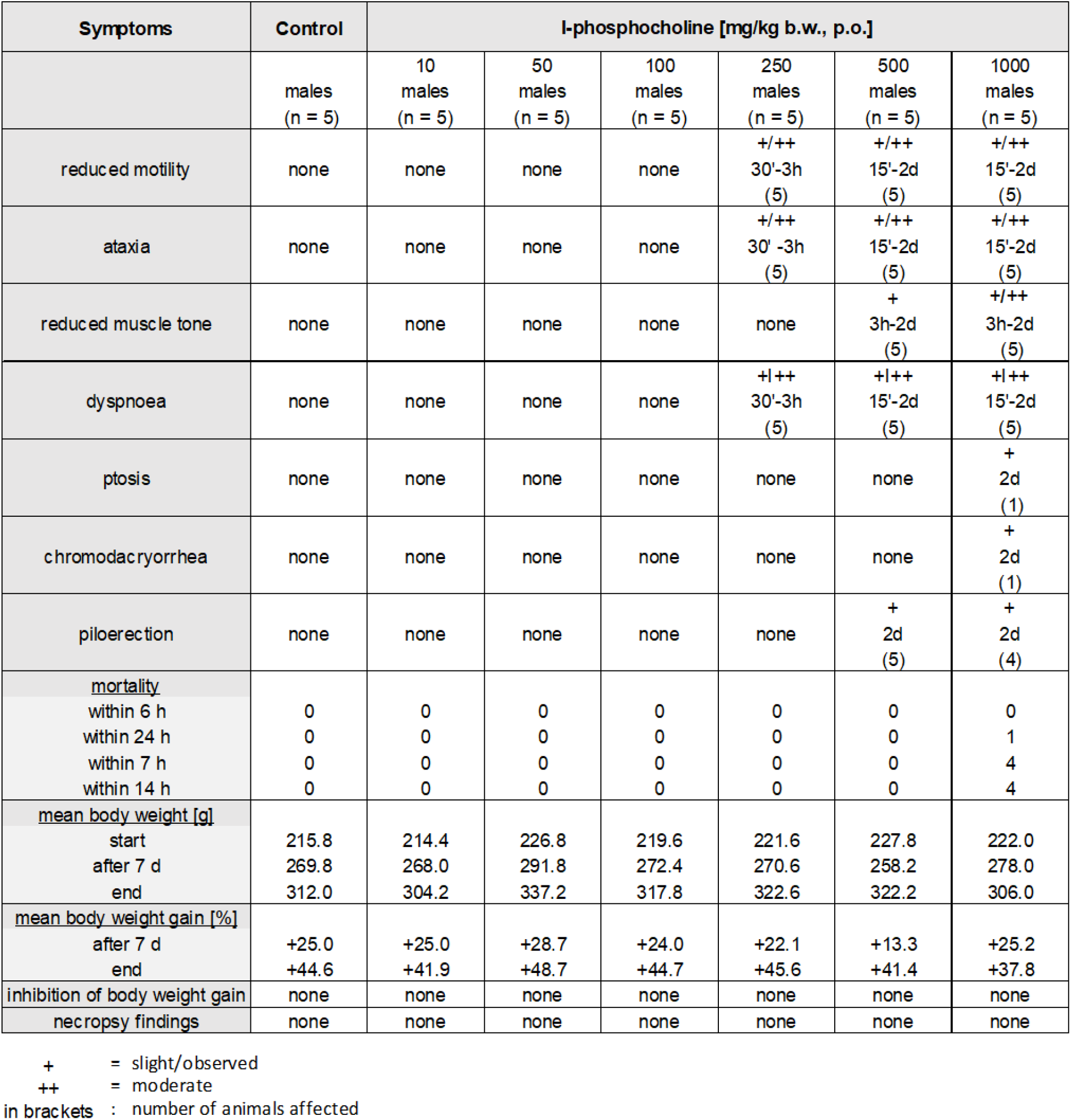
Acute toxicity study results of I-phosphocholine.

